# Temporal probabilistic modeling of bacterial compositions derived from 16S rRNA sequencing

**DOI:** 10.1101/076836

**Authors:** Tarmo Äijö, Christian L. Müller, Richard Bonneau

**Affiliations:** Center for Computational Biology, Simons Foundation, New York, NY 10010, USA; Department of Biology, Center for Genomics and Systems Biology, New York University, New York, NY 10003, USA; Courant Institute of Mathematical Sciences, New York University, New York, NY 10003, USA

## Abstract

The number of microbial and metagenomic studies has increased drastically due to advancements in next-generation sequencing-based measurement techniques. Statistical analysis and the validity of conclusions drawn from (time series) 16S rRNA and other metagenomic sequencing data is hampered by the presence of significant amount of noise and missing data (sampling zeros). Accounting uncertainty in microbiome data is often challenging due to the difficulty of obtaining biological replicates. Additionally, the compositional nature of current amplicon and metagenomic data differs from many other biological data types adding another challenge to the data analysis.

To address these challenges in human microbiome research, we introduce a novel probabilistic approach to explicitly model overdispersion and sampling zeros by considering the temporal correlation between nearby time points using Gaussian Processes. The proposed Temporal Gaussian Process Model for Compositional Data Analysis (TGP-CODA) shows superior modeling performance compared to commonly used Dirichlet-multinomial, multinomial, and non-parametric regression models on real and synthetic data. We demonstrate that the nonreplicative nature of human gut microbiota studies can be partially overcome by our method with proper experimental design of dense temporal sampling. We also show that different modeling approaches have a strong impact on ecological interpretation of the data, such as stationarity, persistence, and environmental noise models.

A Stan implementation of the proposed method is available under MIT license at https://github.com/tare/GPMicrobiome.

## 1 Introduction

Microbial ecology involves the study of microorganisms’ relationships with each other and with their environment and aims to provide insights into structure and dynamics of ecological networks (Kurtz *et al.*, 2015), ecological stability (Faith *et al.*, 2013), biodiversity (Lozupone *et al.*, 2012), and discovery of key taxa in ecosystems (Ivanov *et al.*, 2009).

16S ribosomal RNA (rRNA) amplicon sequencing (targeted next-generation sequencing of 16S rRNA gene) has proven to be a cost-effective, culture-free, and highly multiplexed method to identify and compare bacterial compositions present within biological samples across a wide range of habitats, including natural environments (Meron *et al.*, 2012; Hell *et al.*, 2013) and different host organisms (Kuczynski *et al.*, 2012; Yatsunenko *et al.*, 2012). While the majority of amplicon sequencing studies has been cross-sectional in nature or based on few selected time points, it has been recognized that longitudinal studies with the aim of mapping the trajectories of microbiota over time are a prerequisite for a deeper understanding of ecological mechanisms in the microbiome and for the development of microbiome therapies (Gerber, 2014; Fisher and Mehta, 2014). Sparsely sampled microbial time series have already revealed dynamic reorganization of gut microbial compositions during early development in humans (Yatsunenko *et al.*, 2012) and upon external perturbations through antibiotic treatment (Jernberg *et al.*, 2010), and have identified significant differences in vaginal microbiota during pregnancy (Romero *et al.*, 2014). The richest resource to date for long-term longitudinal amplicon studies are the landmark studies by *Caporaso et al.* (2011) and *David et al.* (2014) which provide human-associated microbial compositions on a daily time scale spanning hundreds of days. *Caporaso et al.* (2011) quantify natural variations of microbial compositions within and among four body sites across time. David *et al.* (2014) focus on the effects of host lifestyle, including travel, change of diet, and infection, on changes in the human gut microbiome.

While statistical time series analysis has an extensive and successful history in classical genomics (Aach and Church, 2001; Bar-Joseph *et al.*, 2004; Bonneau *et al.*, 2006; Leek *et al.*, 2006; Ahdesmäki *et al.*, 2007; Bar-Joseph *et al.*, 2012; Äijö *et al.*, 2014), few attempts have been made to model amplicon-based temporal data in a principled statistical manner (Gerber *et al.*, 2012; Bucci *et al.*, 2016). This may stem in part from the fact that standard multivariate techniques can not be applied to amplicon-based sequencing data. Firstly, as compared to other technologies such as flow cytometry (Amann *et al.*, 1990) and conventional plate counting that allow absolute taxa abundance measurements, standard 16S rRNA count data can only reveal *relative* abundances of taxa, thus rendering individual taxa counts not independent. Secondly, statistical analysis of 16S rRNA sequencing count data is complicated by the presence of overdispersion and missing data. Missing data manifests as an excessive number of zero counts due to imperfect sampling (i.e, zero-inflation and sampling zeros). Separation of sampling zeros (zeros due imperfect sampling) from structural zeros (true, biologically meaningful, zeros) is a common challenge in the analysis of many current biological data types, including single-cell RNA sequencing (Brennecke *et al.*, 2013) and shotgun protein mass spectrometry data (Webb-Robertson *et al.*, 2015). In the context of human-associated microbiome studies, amplicon-based sequencing studies face the additional restriction that well-controlled biological replicates (from different individuals) are not available due to different genetic background, environmental exposure, and life style of human subjects.

Different approaches have been proposed to deal with these intrinsic characteristics of (cross-sectional) 16S rRNA sequencing data (see, e.g., *Xu et al.* (2015) for a recent comparison). Methods based on the negative binomial (NB) distribution (popular in modeling RNA sequencing data) have been proposed for modeling overdispersion in 16S rRNA data, and zero-inflated negative binomial (ZINB) mixture models have been successfully used to fit excessive numbers of zeros. However, the NB and ZINB distributions model taxa as independent, thus ignoring the intrinsic compositional nature of the data. Moreover, the binary distribution component of ZINB only increases the probability of zeros instead of modeling the source of zeros (true vs. non-detected due to sequencing depth) (Mohri and Roark, 2005). The impossibility of obtaining well-controlled biological replicates of human microbiome samples limits the applicability of NB distribution and ZINB in that context because overdispersion of (taxon-specific) counts caused by biological variation cannot be reliably estimated. In light of these limitations, several methodologies have been proposed for simultaneous modeling of taxa through their relative abundances, such as the Dirichlet-multinomial (DM) (Holmes *et al.*, 2012; Chen and Li, 2013) and logistic normal multinomial models (Xia *et al.*, 2013). The logistic normal multinomial model is a generalized linear model (GLM) utilizing the logit link function, thus enabling the use of well-established theory and methods of linear models for modeling count data and relative abundances. Both models are extremely powerful for cross-sectional studies with proper biological replicates. Yet, extending these models to time course data analysis has thus far been limited to point-wise analysis, followed by projecting the dynamics using low-dimensional embedding (Caporaso *et al.*, 2011) or calculating different diversity metrics or temporal summary statistics across pairs of time points (Flores *et al.*, 2014; Faust *et al.*, 2015). Recent approaches that utilize the full potential of the data by considering temporal dependencies among the data points include MC-TIMME (Gerber *et al.*, 2012) which uses exponential relaxation processes to model time-varying counts (Gerber *et al.*, 2012) and BioMiCo (Shafiei *et al.*, 2015) which uses a supervised hierarchical mixed-membership model to track groups of taxa over time. Other methods rely on deterministic regularized model fitting using generalized Lotka-Volterra equations (Stein *et al.*, 2013; Buffie *et al.*, 2015; Bucci *et al.*, 2016).

In this study, we present a fully Bayesian probabilistic model, the Temporal Gaussian Process Model for Compositional Data Analysis (TGP-CODA), that tackles the compositionality, overdispersion, and zero-inflation in 16S rRNA sequencing data through temporal analysis. Our approach is based on the assumption that by sharing information across time points it is possible to improve inference of overdispersion and zero-inflation parameters. We demonstrate that our model can accurately distinguish sampling zeros from structural zeros by using the temporal correlation and the global effect of sampling zeros on the compositions. Our generative hierarchical model combines a multinomial distribution with Gaussian processes (for each taxon to model connections between time points), includes explicit model-based zero-inflation and overdispersion components, and can seamlessly integrate non-uniformly sampled time series (Section 2). We compare our temporal approach to the state-of-the-art DM model on realistic synthetic data and demonstrate more accurate composition estimation. We also model and reanalyze the long-term longitudinal gut microbiota data sets of four individuals (Caporaso *et al.*, 2011; David *et al.*, 2014) using TGP-CODA and maximum likelihood approaches (Section 3). We demonstrate (1) that the dynamical behavior of bacterial orders are globally stable but can accelerate upon environmental perturbations, (2) that our Bayesian model is robust to missing time points, and (3) that estimates of fundamental ecological indicators such as taxa persistence times and taxa stationarity are dependent on the underlying temporal model.

## 2 Methods

We first describe TGP-CODA, our Bayesian generative model that integrates temporal, overdispersion, and zero-inflation components for analyzing longitudinal 16S rRNA sequencing data (Figure 1).

**Figure 1:**
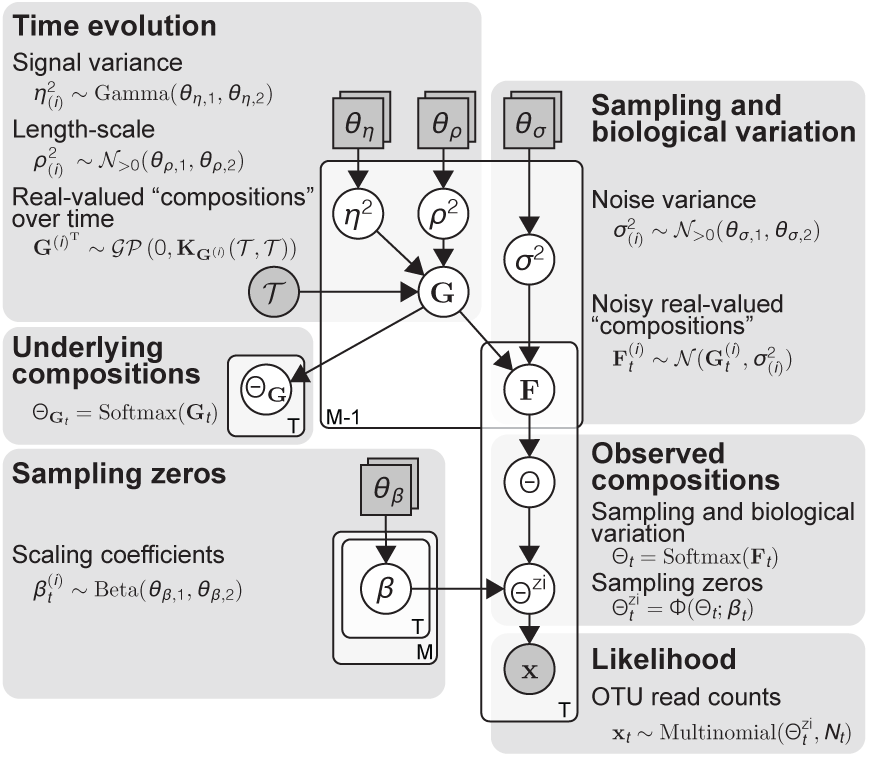
Statistical model and prior distributions. A graphical representation of our model. Grey and white circles depict observed variables and latent variables, respectively. Grey squares represent user-definable parameters. The Gaussian processes, **G**, model noise-free real-valued “compositions” (log odds ratios), which are used as a basis for generating noisy real-valued “compositions” (log odds ratios), **F**. Noisy compositions, Θ, are obtained from **F** by applying the softmax transformation. Zero-inflation-aware compositions, Θ^zi^, are obtained from Θ and *β* by Θ^zi^ = Φ(Θ; *β*) (Equation (13)). The likelihood of data is evaluated using the zero-inflation-aware composition parameters, Θ^zi^. Underlying unobservable noise-free compositions, Θ_**G**_, are obtained from **G** by applying the softmax transformation.

### 2.1 Data likelihood

Let *M* be the number of taxa, *T* the number of measurement time points, and 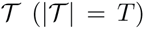 the set of measurement time points. Let 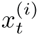 be the number of observed reads assigned to the *i*^th^ taxon at time point 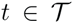 (the corresponding random variable is denoted by 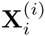, where every read is assigned exactly to one taxon. For notational simplicity, let 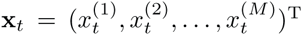 and 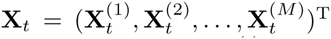. Additionally, let us denote the total number of taxa assigned reads at time point *t* by 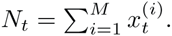. Next, let us assume: (1) *N*_*t*_ taxa reads are sampled independently of each other and (2) the *M* possible outcomes have fixed probabilities, 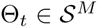 (*M*-dimensional simplex), at time point t. Then, **X**_*t*_ follows multinomial distribution with the parameters Θ_*t*_ and *N*_*t*_

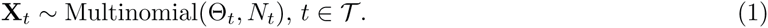

The normal approximation to the multinomial (Severini, 2005), while computationally convenient, is not applicable in this case even for large values *N*_*t*_ because Θ_*t*_ is empirically observed to be located close to a corner of the simplex 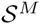(i.e., there are many lowly abundant taxa).

Next, let us define the likelihood in the case of multiple time points. Let us denote the collection of Θ_*t*_ over *T* time points by:

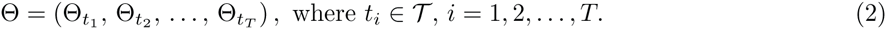

The data likelihood assuming independence of observations at different time points (true for sequential sampling from a population) (Figure 1; see the “Likelihood” section), 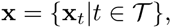, is

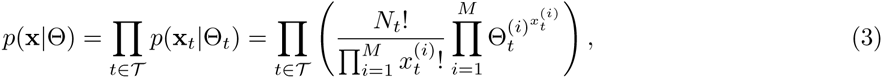

which can be used to evaluate the likelihood of the data, 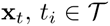 given the parameter Θ_*t*_.

### 2.2 Temporal modeling of microbiome compositions

Modeling in compositional space is notoriously challenging (modeling fractions of population or fractions of reads, for example) (Aitchison, 1982): (1) the compositional space enforces restrictions on the modeling domain, which might not be easily expressible in the selected modeling framework (due to the intrinsic dependency among all taxa) and (2) the differences in relative abundances of taxa can vary over multiple orders of magnitude, which, combined with compositional effects renders the direct modeling of relative abundances a hard task. To overcome these challenges, modeling log odds ratios between taxa in real space have been proposed, typically followed by a transformation to map the real values to a simplex (Aitchison, 1982; Holmes *et al.*, 2012). In this study, we will use the commonly used softmax transformation (e.g., in multinomial logistic regression) which is a generalization of the logistic function (Bishop, 2006). The softmax transformation from 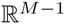 to 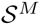 is defined as follows

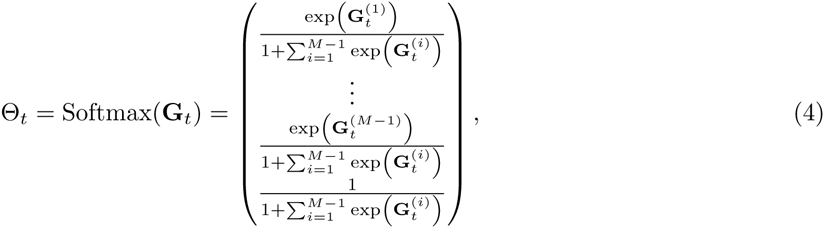

where 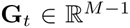(Bishop, 2006). The explicit assumption 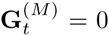 in Equation (4) makes the softmax transformation bijective. The softmax transformation is required because the multinomial likelihood parameters, Θ_*t*_, are constrained to lie in the M-dimensional simplex. Next, let us denote the collection of **G**_*t*_ over *T* time points by

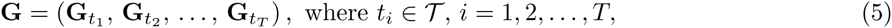

with the element-wise softmax transformation (see also Equation (2))

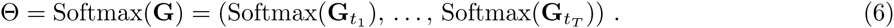

Next, we will describe the temporal component of our generative model. It is unknown *a priori* how relative abundances of bacterial taxa vary over time and how treatments and abrupt changes in the environment might alter ecological dynamics. Therefore, we do not want to restrict the model and the resulting dynamics by strong assumptions on functional forms of temporal relative abundances. Thus, we will take a non-parametric approach and use a Gaussian process kernel to model temporal dynamics, requiring only weak assumptions (such as smoothness) on the temporal characteristics of the signal (Rasmussen and Williams, 2005).

We assume that **G**^(*i*)^,*i* = 1,2,…,*M* − 1 (*i*^th^ row of **G**) are smooth, and the time series data is well sampled (i.e., well-designed experiments to match the modeling objective). We will model **G**^(*i*)^,*i* = 1,2,…,*M* − 1 using Gaussian process (Rasmussen and Williams, 2005)

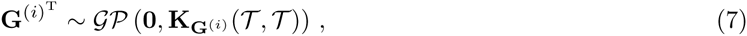

where 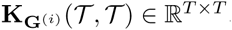, *i* = 1,2,…,*M* − 1 using Gaussian process (Rasmussen and Williams, 2005)

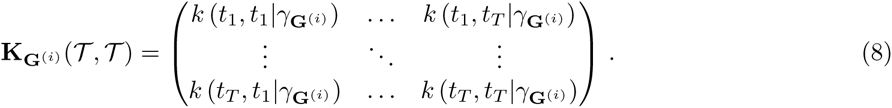

The term 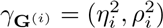 is the covariance function given the hyperparameters 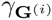. In this work, we use the squared exponential covariance function

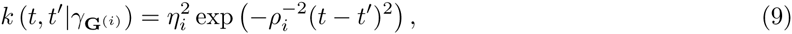

where 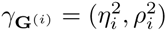 with *η*_*i*_ denoting the signal variance parameter and *ρ*_*i*_ the characteristic length scale.

### 2.3 Modeling overdispersion of counts

When the values *N*_*t*_ are large and no replicates are available, the data likelihood (Equation (3)) will dominate the Gaussian process prior (Equation (7)) leading to overfitting of Θ_*t*_. Consequently, inherent biological and technical variations are severely underestimated. Notably, the DM and logistic normal multinomial models suffer from the same problem (this is apparent from the forms of maximum likelihood and Bayes estimators in SEquations (1) and (4), respectively). Thus, it is advantageous to explicitly model sampling variation in 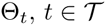 by introducing an additional level of random variables to the hierarchical model

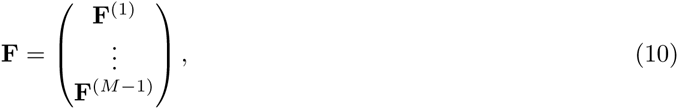

where 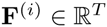, *i* = 1,2,…,*M* − 1 are row vectors that depend on **G**^(*i*)^ and 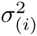 as follows:

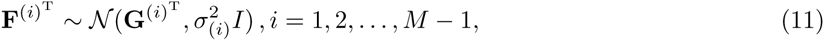

where 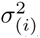 is assumed to be constant over time (i.e., sampling variation is similar over time series) in order improve identifiability. In this extended model, Θ is obtained by applying the softmax transformation on **F**(see also Equation (2))

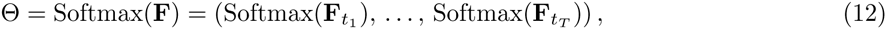

where 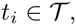, *i* = 1, 2,…, *T*.

In summary, the random variable Θ = Softmax(**F**) (see Equations (11) and (12)) is sample-specific (after sampling), whereas the random variable Θ_**G**_ = Softmax(**G**) (see Equations (7) and (6)) models biological variation over samples (before sampling). The overdispersion component of the model is illustrated in Figure 1 (see the “Sampling and biological variation” and “Observed compositions” sections).

### 2.4 Modeling zero-inflation and missing data

16S rRNA and other amplicon sequencing based count data have been empirically shown to suffer from severe zero-inflation (Xu *et al.*, 2015). Zero-inflation can be seen as “salt” noise in the compositions Θ_*t*_ (i.e., zeroing of individual components of Θ_*t*_); the “salt” term refers to the “salt-and-pepper” noise concept from the digital image processing literature (Jayaraman, 2009). To model zero-inflation, we introduce another level of simplex-valued latent variables, 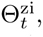 to the model (Figure 1). The variables Θ_*t*_ and 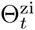 model underlying proportions and “salty” proportions of taxa, respectively. The sampling and zero-inflation are modeled separately for modeling convenience and for identifying the *source* of zeros (sampling or structural).

To explicitly model the effect of imperfect sampling, we introduce random variables 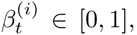,*i* = 1,2,…,*M* and consider the following weighting based transformation:

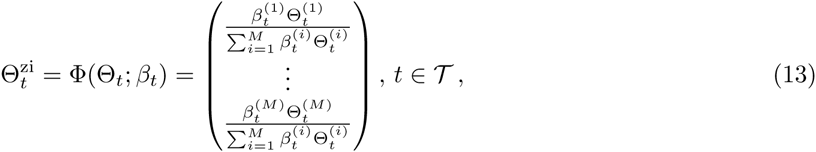

where the common denominator term ensures 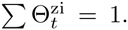. For notational simplicity, let us denote *β*=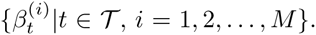. The zero-inflation component of the model is illustrated in Figure 1 (see the “Sampling zeros” and “Observed compositions” sections).

### 2.5 Posterior estimation

To carry out the Bayesian inference on the presented model (Figure 1), we first specify the parameter prior distributions, 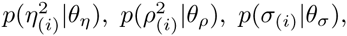 and 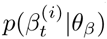 (SFigure 1a). The parameters 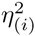 and 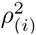 determine the signal variance and how fast correlation between time points diminishes, respectively. We select a relatively broad prior distribution for 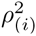 in order to support temporal correlations that vary from a few days to a few weeks (SFigure 1a). In this study, the time points *t*_*i*_ (model inputs) are obtained by scaling the days of measurement (e.g., integers from 1 to D) by the total number of days (D); thus, the prior of 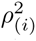 is selected as 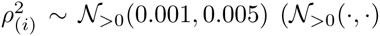 is positive truncated Gaussian distribution) (SFigure 1a). Since Gaussian processes model the log odds ratios, we assume that the variances of the log odds ratios of taxa over time are relatively small. We set the prior as 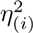 ~ Gamma(1.0, 0.5) (SFigure 1b). The prior of the noise standard deviation is set to 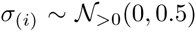 to support relatively low noise levels (SFigure 1b). Finally, we explicitly assume that the sampling zeros are relatively rare by defining the prior as 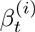 ~ Beta(0.8, 0.4) (broad distribution improves sampling efficiency) (SFigure 1b).

The posterior distribution function (up to a normalizing constant) is obtained as the product of the likelihood function and priors. The full posterior distribution function of our model is given in SEquation (5). We implemented the model in Stan (Carpenter *et al.*, in press) and used its No-U-Turn Sampler (NUTS) to sample the posterior (SEquation (5)). The Stan probabilistic programming language enables cross-platform implementation, code interpretability, numerical stability, scaling, and efficient posterior inferences of various statistical models. Convergence of chains was monitored using by the Gelman-Rubin statistic (Gelman and Rubin, 1992) 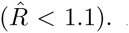. All relevant information (prior and data) about the parameters is summarized in the posterior distributions. We can thus use the obtained posterior samples to summarize the distributions, e.g., by calculating means and credible intervals (Gelman *et al.*, 2014).

## 3 Results

### 3.1 Temporal analysis improves estimation accuracy

To validate the presented temporal compositional data analysis method, we first compare TGP-CODA to the DM model (Chen and Li, 2013) using synthetic data. To compare these two methods, we consider a scenario of 36 taxa with realistic dynamics and abundance distribution (see Supplementary Material). The generated synthetic data sets are analyzed using the temporal and DM models. The composition estimates at day 90 (common between six, nine, 14, and 27 time points to allow direct comparison) of both methods are compared to the noise-free ground truths (Figure 2a). Even in this simple scenario, the temporal approach consistently produces more accurate composition estimates than the DM model (Figure 2; STable 1). We find that the performance of the temporal approach improves (as expected) as the number of time points increases; e.g., the mean estimation errors and the corresponding standard deviations are 0.15±0.09 and 0.10±0.06 with six and 14 time points, respectively (STable 1). The estimation error of the DM model does not depend on the number of time points as it considers time points separately (STable 1). Our modeling of temporal correlations and thereby sharing information between time points leads to more accurate estimation of compositions from longitudinal count data.

**Figure 2:**
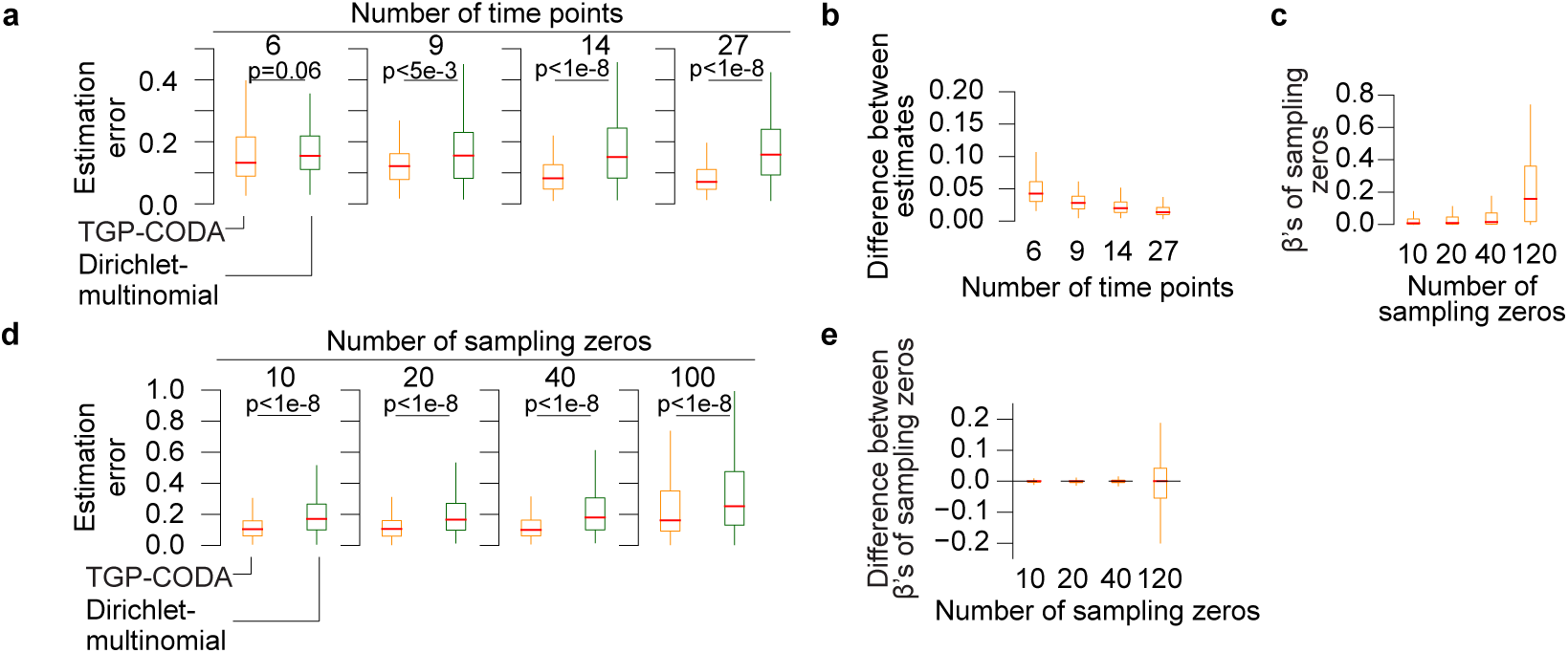
Temporal correlation in composition estimation. (**a**) Box plots illustrate estimation errors of our temporal TGP-CODA (orange) and DM models (green). Six, nine, 14, and 27 time points are considered. Estimation error is defined to be the Euclidean distance between the the first M 1 components of the simplex-valued proportions vectors. Each box plot is calculated from 100 simulations. Outliers are not shown. The two-sided p-values from the Wilcoxon signed-rank tests are listed. (**b**) The results from the sensitivity analysis of the results depicted in (**a**) with respect to the prior distributions of η^2^ and ρ^2^ are illustrated. (**c**) Box plots illustrate the variation of *β* values of taxa (proportions ≥ 1e-4) and time points with added sampling zero. The cases of 14 time points with either 10, 20, 40, or 120 added sampling zeros are considered. Each box plot shows the average from 100 simulations. Outliers are not depicted. (**d**)Box plots illustrate the estimation error of the temporal (orange) and DM (green) models at the time points with induced sampling zeros. The cases of 10, 20, 40, and 120 sampling zeros are considered. Estimation error is defined to be the Euclidean distance between the the first M-1 of the simplex-valued proportions vectors. Each box plot is calculated from 100 simulations. Outliers are not depicted. The two-sided p-values from the Wilcoxon signed-rank tests are listed. (**e**) The sensitivity of the estimation of *β* with respect to the prior of *β* is studied in the scenario of (**c**). The estimated *β* values of taxa (proportions ≥ 1e-4) and time points with induced sampling zeros are compared by calculating the difference between the estimates obtained under the original prior (*β* ~ Beta(0.8, 0.4)) and perturbed prior (*β* ~ Beta(θ_*β*,1_,*θ*_*β*,2_)). The cases of 14 time points with either 10, 20, 40, or 120 sampling zeros are considered. Each box plot is calculated from 100 simulations. Outliers are not depicted.

Because our estimates should not be critically sensitive to the hyperparameters, (*θ*_*η*_, *θ*_*ρ*_, *θ*_*β*_). we carried out a sensitivity analysis with respect to the prior distributions of *η*^2^ and *ρ*^2^ defined in the Section 2.5. We considered random variables 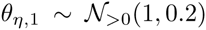 and 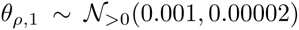(SFigure 4a) whose purpose is to perturb the prior distributions 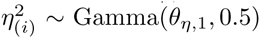 and 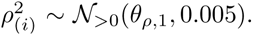. We then repeated the analysis presented in Figure 2a and compared the compositions estimates between the original and perturbed priors (Figure 2b). The means and the corresponding standard deviations of the estimate differences were 0.05±0.05, 0.03±0.03, 0.03±0.08, and 0.02±0.01 with six, nine, 14, and 27 time points, respectively (Figure 2b). As expected, the variations in the final estimates get smaller as the amount of data to base the estimation increases. Collectively, the small obtained differences demonstrate that the estimates are not critically sensitive to the prior distributions of η^2^ and ρ^2^.

### 3.2 Modeling sampling zeros improves estimation accuracy

To validate the described zero-inflation component and to see whether the estimated β values reflect sampling zeros, we consider the same example as above but with imposed sampling zeros. We generated data sets with different numbers (10, 20, 40, or 120) of imposed zeros randomly distributed to the taxa and time points. Importantly, there are likely additional zeros for lowly abundant taxa due to the low sampling depth. This unbiased procedure also introduces sampling zeros to lowly abundant taxa. Clearly, these zeros are harder to detect with the used sampling depth (or with any relatively low sampling depth). We analyzed these zero-inflated synthetic data sets using our temporal approach and studied the distribution of *β* values of taxa (proportions ≥ 1e-4) at the time points with imposed zeros (Figure 2c). Our model is able to identify 10 (mean±SD=0.04±0.07), 20 (0.05±0.09), and 40 (0.07±0.11) sampling zeros accurately among taxa that are not close to detection limit, whereas the identification of 120 sampling zeros is less reliable (0.21±0.19) (Figure 2c). As expected, detecting sampling zeros among lowly abundant taxa is challenging (SFigure 4b).

To check whether the detection and correction of sampling zeros improves composition estimation, we next considered the same scenario as shown in Figure 2c with the focus on the composition estimates instead of *β* value. We compared the composition estimates of the temporal and DM models at the time points with sampling zeros to the noise-free ground truths (Figure 2d, STable 2). The temporal approach produces smaller estimation error than the DM model in all the considered cases. For instance, the estimation error is almost two times smaller with the temporal approach (mean*±*SD=0.12*±*0.08) compared to the DM model (0.21±0.16) in the case of 20 sampling zeros (Figure 2d, STable 2). The weaker performance of the DM model is expected since it does not explicitly model sampling zeros. Additionally, we repeated this analysis with greater numbers of taxa (71, 102, and 160) and sampling zeros (120, 240, and 480) and 27 time points to validate our model’s performance in a larger setting (SFigure 4d).

To confirm that the estimation of sampling zeros is not critically sensitive to the prior distribution of *β*, we considered a perturbed prior, *β* ~ Beta(*θ*_*β,1*_, *θ*_*β,2*_) where *θ*_*β,1*_ ~ Beta(16, 4) and *θ*_*β,2*_ ~ Beta(8, 12) (SFigure 4c). Then, we compared the *β* estimates obtained with the original and perturbed prior in the case of Figure 2c (Figure 2e). The *β* estimates were stable with respect to the prior distribution; the means and the corresponding standard deviations of the differences were -0.006*±*0.044, 0.006*±*0.056, -0.005*±*0.066, and -0.011*±*0.134 with 10, 20, 40, and 120 sampling zeros, respectively.

### 3.3 Differential response of bacterial orders to environmental perturbations

To demonstrate our approach on real data, we reanalyzed the longitudinal gut microbial 16S rRNA sequencing data sets of four individuals, referred to as M3 and F4 (Caporaso *et al.*, 2011) and Subject A and B (Caporaso *et al.*, 2011; David *et al.*, 2014) (see Supplementary Material). The percentages of zeros varied between 66% and 78% in these data sets (see Supplementary Material). Due to the sparsity of the data we grouped all Operational Taxonomic Units (OTUs) according to phylogenetic order and analyzed the resulting compositions. We visualize the dynamics of the orders in SFigures 5–8 by plotting the posterior mean composition estimates of bacterial orders with corresponding credible intervals at time points with and without measurements. For comparison, we included the maximum likelihood estimates (MLEs) under the multinomial model with and without the locally weighted scatterplot smoothing (LOWESS) (Cleveland, 1981).

We first focus on the Subject B time series. From days 151 to 159 the subject had a Salmonella infection; as expected, relative abundance of Enterobacteriales increases upon the infection as reported in (David *et al.*, 2014) (SFigure 6a). Similarly, relative abundance of Enterobacteriales in Subject A’s gut microbiota is greater during the travel abroad (SFigure 5a). The relative abundance of Bifidobacteriales decreases during the time Subject A spent abroad (from 7e-2 to 2e-2) (SFigure 5a). The disappearance of the RF39 order from the gut microbiota of Subject B coincides with the Salmonella infection (average relative abundances pre-infection and post-infection are 5e-3 and 8e-7, respectively) (SFigure 6a). The decrease in the relative abundance of Enterobacteriales in F4’s gut microbiota around 50 days coincides with the increase of the relative abundances of Burkholderiales (SFigure 8a). Interestingly, our results suggest that F4’s gut microbiota undergo a global transition between states around 50 days (SFigure 9). Identification of the importance and/or the cause of this would require additional metadata. Finally, TGP-CODA quantifies the uncertainty in estimates caused by lower sequencing depth and missing samples (e.g., see lowly abundant orders Gallionellales in SFigure 5, Acidimicrobiales in SFigure 6, and Gammaproteobacteria in SFigure 8).

To confirm that the results are not too sensitive to the selected covariance function, we reanalysed the Subject A data using the Matérn covariance function (*v* = 3/2). The obtained similar results suggest that our method is stable with respect to the chosen covariance function (SFigures 9,10); the slightly less smooth processes are expected as the Matérn covariance function (with *v* = 3/2) leads to processes that are 1-times mean square (MS) differentiable, whereas the squared exponential covariance function leads to processes that are infinitely MS-differentiable. Additionally, to verify that our method does not produce analysis artifacts due to the temporal modeling, we shuffled the time points in the Subject A data set and analyzed the shuffled data (SFigure 11). As expected, we did lose the signals observed with the original data (SFigure 9). Importantly, the LOWESS estimator does seem to overfit the shuffled data (SFigure 11).

Collectively, our temporal approach is able to recover patterns from highly noisy 16S rRNA data which are not apparent from the MLEs even when these perturbations effect extreme restructuring of the dynamics and composition of the niche.

### 3.4 Effect of sampling frequency on estimating microbiome dynamics

To see study how the data sampling frequency affects the results, we performed downsampling experiments. Specifically, we reanalyzed Subject A data by taking into account only measurements from either every second or third time point (SFigures 12,13). Overall, the obtained results with the full and downsampled data sets are highly similar suggesting that daily sampling is not necessary to capture human gut microbiota dynamics (SFigures 4,9,12,13). In Figure 3, we illustrate four examples of how different sampling frequencies can affect results. As expected, when sampling frequency drops credible intervals become wider (see Bacteroidales in Figure 3). Importantly, the LOWESS estimates are sensitive to the sampling frequency, which suggests that the LOWESS estimator tends to overfit data (see Enterobacteriales, Sphingomonadales, and Myxococcales in Figure 3). The observed overfitting, especially among lowly abundant orders, is not surprising since LOWESS and ML estimation do not take into account the statistical nature of count data. Additionally, in contrast to our method, it is not straightforward to interpolate data with LOWESS.

**Figure 3:**
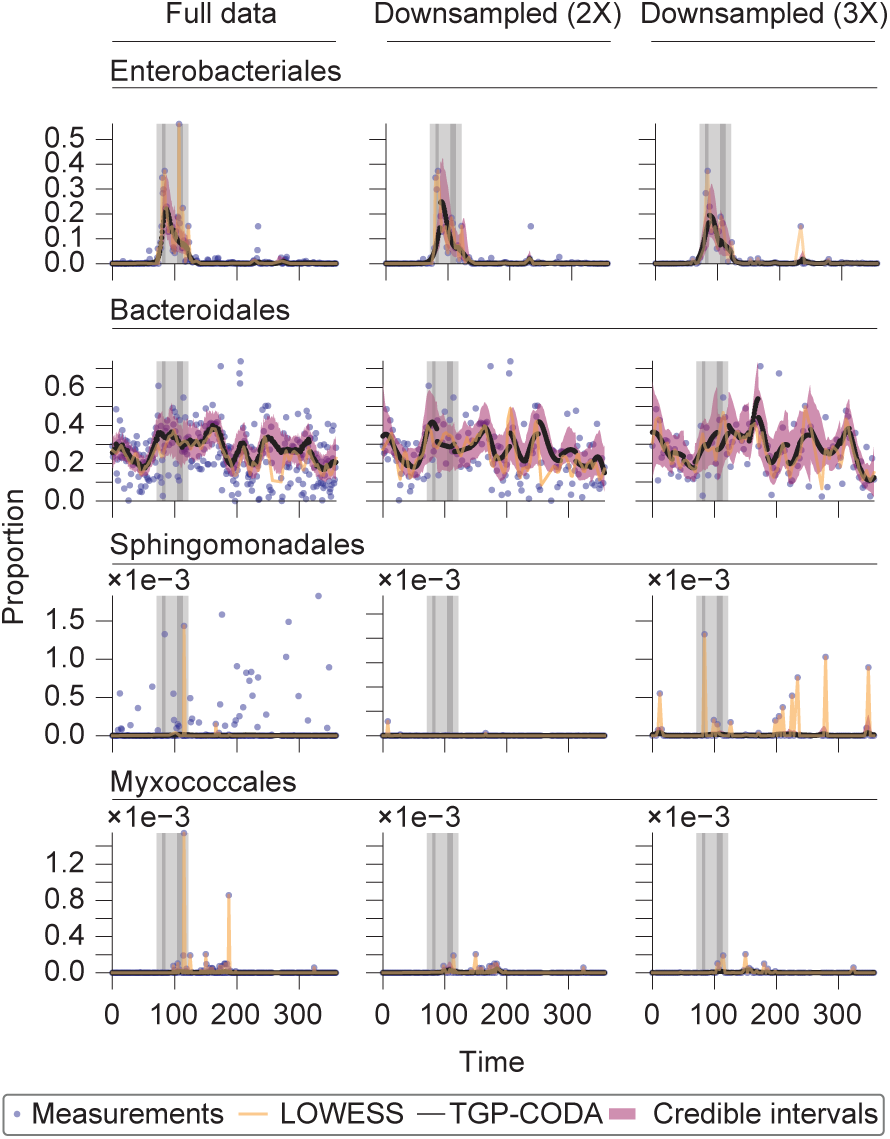
Effect of sampling frequency on the estimation of bacterial order dynamics. (**a**) Dynamics of the proportions of Enterobacteriales (first row), Bacteroidales (second row), Sphingomonadales (third row), and Myxococcales (fourth row) in Subject A’s gut microbiota over time. The black circles are the posterior mean estimates, Θ_**G**_, from the temporal analysis. The filled regions show the 5% and 95% credible intervals. The semi-transparent circles depict the maximum likelihood estimates under the multinomial model. The orange curve is the LOWESS (*α* = 0.05, which corresponds approximately to 20 days) estimate calculated from the maximum likelihood estimates. The time period where the subject was abroad and suffered from diarrhea are illustrated using the three shaded rectangles. (**b**) As in (**a**) but in the case when only every second time point is considered. (**c**) As in (**a**) but in the case when only every third time point is considered.

### 3.5 Revisiting dynamics of human gut microbiota

We next analyze the dynamical properties of the inferred time series and their ecological implications. Our Bayesian framework, together with the use of separate analysis windows, enables us to study the posterior distributions of length scales *ρ*_*i*_ inferred from the different time series. These distributions can serve as global summary statistics of the whole gut microbiota dynamics upon environmental perturbations. We illustrate the results for the Subject A time series over all the bacterial orders in Figure 4a. We first compare the profiles of prior and posterior distributions. We observe that the experimental data supports longer length-scales (i.e., greater temporal correlation) (Figure 4a; see SFigure 1b for interpretation) suggesting that the smoothness of the obtained profiles is not merely an analysis artifact caused by the length-scale prior (SFigure 1a).

**Figure 4:**
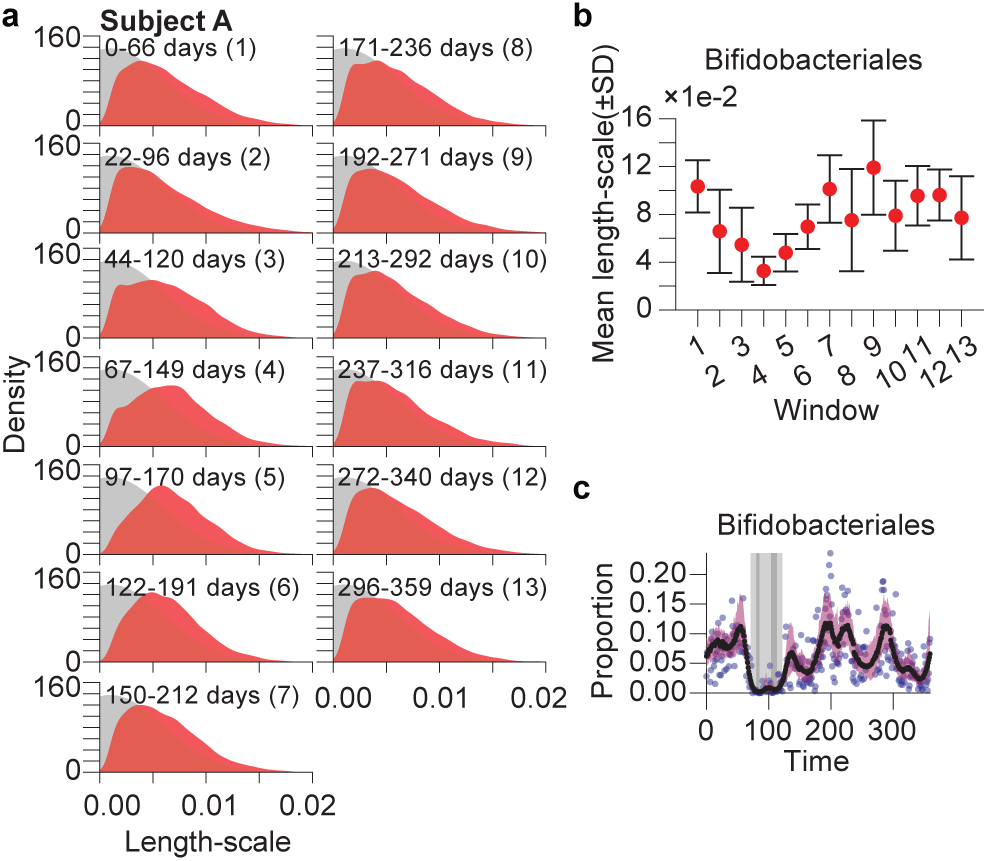
Kinetics of Subject A’s gut microbiota. (**a**) Black and red shaded regions are prior and posterior distributions of the length-scale parameter, respectively. The posterior distributions obtained in different analysis windows are illustrated separately (the days corresponding to each of the windows are listed in the titles). Posterior densities are estimated using Gaussian kernel density estimation (the Scott’s rule for estimating the bandwidth) on the pooled length-scale posterior samples over all the bacterial orders. (**b**) The posterior mean of the length-scale parameter and the corresponding standard deviations of Bifidobacteriales in different analysis windows (the window numbers correspond to the ones listed in (**a**)). (**c**) Dynamics of Bifidobacteriales in Subject A’s gut microbiota over time. The black circles are the posterior mean estimates, Θ_**G**_, from the temporal analysis. The filled regions show the 5% and 95% credible intervals. The semi-transparent circles depict the maximum likelihood estimates under the multinomial model. The time period where the subject was abroad and suffered from diarrhea are illustrated using the three shaded rectangles.

Across all windows, the posterior distributions have an overall similar right-skewed shape and cover a wide range of length scales. This suggests that, on the population level, each bacterial order has different degrees of internal temporal correlations that are persistent across the entire time series (Figure 4). We can also identify several bacterial orders that change their kinetics upon perturbations, as reflected in a potential bi-modality of the distribution between 44 and 149 days (windows 3 and 4 in Figure 4a). To highlight the effect of environmental perturbations, we visualize the length-scale distributions of Bifidobacteriales (Figure 4b,c). The dip in average length scale between windows 2 to 6 suggest that Bifidobacteriales’ kinetics are accelerated upon traveling abroad and being exposed to novel diet.

We next analyze estimates of autocorrelation, persistence, and self-affinity (self-similarity) for the most abundant bacterial orders (mean relative abundance >1e-3 across all four time series under TGP-CODA and ML modeling. We first calculate the sample autocorrelation function (ACF) for lags up to *k* = 60 (SFigure 14a). The TGP-CODA-derived time series show consistently longer autocorrelations (close to 1 in most cases) than the ML-based time series. For most bacterial orders, positive autocorrelation exists for up to a month under TGP-CODA. Coriobacteriales shows particularly strong long-term positive autocorrelation for both Subject A and B. To estimate the degree of self-affinity and the temporal persistence of the bacterial orders we use Hurst’s rescaled range analysis (Hurst, 1951; Di Matteo *et al.*, 2003), resulting in scaling estimates of the Hurst exponent *H* ϵ [0, 1] (SFigure 14b). For ML-based time series we consistently estimate low H values across all time series (mean *H* ϵ[0.15, 0.25]), indicative of memory-less underlying processes, whereas TGP-CODA modeling results in considerably larger Hurst exponent estimates (mean *H* ϵ [0.8, 0.85]), hinting at underlying persistent, self-affine, long-term memory processes. Spectral analysis of the TGP-CODA-modeled times series reveals a scaling of the power spectrum *S(f)* ~ 1/*f* ^*β*^ with *β* ϵ [1.7, 4.2] for the majority of orders (SFigure 15). These results indicate that most time series modeled with TGP-CODA show non-stationary fractional Brownian motion behavior with long-term memory, persistence, and self-affinity.

## 4 Discussion and conclusions

The difficulty of obtaining well-controlled biological replicates renders the estimation of biological and technical variation from individual time points impractical, thus severely limiting interpretability of human microbiome studies. To overcome this limitation, we have derived a probabilistic model, the Temporal Gaussian Process model for Compositional Data Analysis (TGP-CODA), that comprises non-parametric temporal, explicit overdispersion, and zero-inflation noise components leveraging temporal relationships between time points and integrative analysis of all the bacterial taxa (to account for population structure and the compositional nature of typical microbiome data sets). Our results demonstrate that the lack of replicates for longitudinal human gut microbial data can be partially mitigated by our method in the case of proper experimental design: dense time series. Our temporal modeling framework can seamlessly incorporate different experimental designs, such as non-equidistant sampling over time, missing time points, and variable sequencing depth. Our framework also quantifies the uncertainty of the final estimates, which is an important property in integrated microbiome studies, where downstream analysis methods might propagate this error.

Our results on real and synthetic data demonstrate TGP-CODA’s validity and superior performance for analyzing longitudinal microbiome data. Temporal autocorrelation and scaling analysis also revealed that ML and TGP-CODA modeling have a fundamental impact on time series characteristics and their ecological interpretation. ML modeling suggests that the observed time series are stationary and possess short-term memory, driven by white noise. TGP-CODA modeling suggests that relative abundances of microbiota are self-affine, persistent, and possess long-term memory, driven by Brownian noise. Using TGP-CODA, the Hurst exponents of the majority of microbial orders are in remarkable agreement to those of long species abundance time series across the tree of life, including fresh water diatoms (*H* = 0.85) and vertebrates (*H* = 0.77) (Arino and Pimm, 1995). Determining the true underlying dynamics as well as the appropriate environmental noise characteristics will be a key objective for future research because these features will have a major impact on our understanding of species persistence in microbial ecosystems and their potential extinction rates (Sugihara and May, 1990; Cuddington and Yodzis, 1999).

This work also suggests several research questions for future experimental and computational studies. Key objectives are to determine (1) which approximations can be made to the probabilistic model without compromising its validity and (2) how improved temporal analysis can be leveraged to estimate directed, time-varying microbial association networks. A key area of future development will also be the application of TGP-CODA-type methods to mixed experimental designs that include both cross-sectional (perturbation, steady-state) data and time series data. One could envision using time series data to estimate taxon specific zero-inflation parameters that serve as more accurate prior for estimates in cross-sectional data. Another important extension of the model would be the inclusion of the spatial information in a unifying GP modeling framework, which would greatly advance our understanding of microbial ecosystems across space and time.

As the prevalence and public availability of dense time series (including hybrid cross-sectional and time series data) in microbiome research will only increase in the near future, the importance of explicit treatments of microbiome dynamics with models like the one presented herein will likely be instrumental for a deeper understanding of microbial ecosystems.

## Acknowledgements

We acknowledge the computational resources provided by the computing group of Simons Center for Data Analysis.

